# Zebra diel migrations reduce encounter risk with lions over selection for safe habitats

**DOI:** 10.1101/165597

**Authors:** Nicolas Courbin, Andrew J. Loveridge, Hervé Fritz, David W. Macdonald, Rémi Patin, Marion Valeix, Simon Chamaillé-Jammes

## Abstract

Diel migrations (DMs) undertaken by prey to avoid visual predators during the day have been demonstrated in many taxa in aquatic ecosystems. We reveal that zebras in Hwange National Park (Zimbabwe) employ a similar anti-predator strategy. Zebras forage near waterholes during the day but move away from them at sunset. We demonstrate that this DM, occurring over a few km, dramatically reduces their night-time risk of encountering lions, which generally remain close to waterholes. By contrast, zebra changes in night-time selection for vegetation types marginally reduced their risk of encountering lions. This may arise from a trade-off between encounter risk and vulnerability across vegetation types, with zebras favouring low vulnerability once DM has reduced encounter risk. In summary, here we (1) quantify the effect of a predator-induced DM in a terrestrial system on the likelihood of encountering a predator, (2) distinguish the effects of the DM from those related to day/night changes in selection for vegetation types. We discuss how revealing how prey partition their risk between predator encounter risk and habitat-driven vulnerability is likely critical to understand the emergence of anti-predator behavioural strategies.

## INTRODUCTION

Prey species attempt to avoid predators that search for them, leading both predators and prey into a spatial game (Sih 1984, Lima 2002, Sih 2005, Laundré 2010), which has ecological and evolutionary implications for both players (Sih 1998, Flaxman et al. 2011) and sometimes other trophic levels (see Rosenheim 2004, Fortin et al. 2005 for spatial trophic cascade). Several simple one-predator-one-prey models have predicted that the race will revolve around the prey resource patches (Sih 2005). Experiments have confirmed in simple settings that predators often search for habitats rich in prey resources rather than for the prey themselves (Sih 1998, Williams and Flaxman 2012). Prey were found to use the richest but most risky resource patches less than expected under the assumption of an optimization of resource acquisition (Sih 2005, Hammond et al. 2007).

These predictions and experiments ignore, however, that the temporal variations in predation risk affect the spatial behaviour of prey. Variations in predation risk may occur at different temporal scales (e.g. day, seasons and year), but it is most obvious when looking at how most predator-prey interactions are affected by the day/night cycle. The hunting efficiency of many predators varies with light intensity, leading many predators to have well defined and restricted windows of hunting activity over the 24h cycle (Clark and Levy 1988, Lima and Dill 1990, Kronfeld-Schor and Dayan 2003). Prey may thus optimize how they balance forage and predation risk by making strategic use of rich resource areas when the predator is inefficient and/or inactive. Thus, during the low-risk period, prey could tolerate predator presence. During the high-risk period, prey could reduce their overall predation risk by decreasing the probability of encountering the predator by moving away from the rich resource areas.

Such spatio-temporal strategy of prey is observed in diel migrations conducted by a wide range of taxa in aquatic ecosystems (Alonzo et al. 2003, Hays 2003, Benoit-Bird and Au 2006). For example, in marine systems, zooplankton forage on phytoplankton at the sea surface at night when their predators have a reduced visual acuity, and move towards deeper water during the day to reduce the risk of being detected, leading to the emergence of diel vertical migrations (Iwasa 1982, Hays 2003). Similarly, diel horizontal migrations have also been reported. For instance, in shallow lakes, zooplankton migrates to the safer vegetated littoral zone during daytime to avoid visual predators (Burks et al. 2002). Thus, diel migrations (DMs) are a common proactive strategy employed by aquatic organisms to exploit their environments in the context of food-safety trade-offs (Hays 2003), while DMs have been largely overlooked for terrestrial prey.

Predation risk arises not only from the risk of encountering the predator but also from the vulnerability of the prey (i.e. the likelihood of dying if attacked) (Prins and Iason 1989, Lima and Dill 1990, Hebblewhite et al. 2005). Therefore, during the high-risk diel period, prey could also remain near the resource rich areas and the predator, but shift to neighbouring safer habitats (Schmidt and Kuijper 2015). In many systems, the vegetation cover, which may change abruptly over short distances, is a strong determinant of predator hunting efficiency (Mysterud and Østbye 1999, Hopcraft et al. 2005) and hence of prey vulnerability. This may for instance be linked to the higher visibility, or the ambush opportunities, that some habitats provide (Lima and Dill 1990, Caro 2005). During diel periods of predator activity, prey can thus decrease predation risk without much travel by shifting their habitat selection towards neighbouring habitat where they are less vulnerable. This is commonly observed in many ungulate species facing natural predators such as wolves (Creel et al. 2005, Middleton et al. 2013, Basille et al. 2015, Schmidt and Kuijper 2015) and puma (Laundré 2010), or hunters (Padié et al. 2015).

The relative efficiency of the two strategies (DM vs. local habitat shift) will likely depend upon the correlation between encounter risk and vulnerability across habitat types and the predator behaviour. If some nearby habitat simultaneously offers lower encounter risk and lower vulnerability, i.e. refuge habitat (Sih 1984, Hays 2003), these should be selected for by prey when the predator is active. If encounter risk and vulnerability are negatively correlated, then there is no refuge habitat and the cost of changing habitat will likely depend on the actual encounter risk and vulnerability within and across habitats and the predator behaviour. Some predators are spatio-temporally unpredictable, either because they roam over large areas in the quest of vulnerable prey (Latombe et al. 2014) or track prey resource patches in landscapes where these are common and scattered (Courbin et al. 2014). The use of habitat shift strategies could then be more efficient to decrease predation risk than a spatial redistribution towards areas for which information on the recent predator activity is not available or not reliable (Creel et al. 2005, Middleton et al. 2013, Basille et al. 2015, Schmidt and Kuijper 2015). Conversely, some predators may be more predictable, anchored near scarce prey resource patches (Sih 2005), relying more on prey attraction to the patch than on selectively tracking individual prey to encounter them (Valeix et al. 2010). In such a context, a DM could be a more efficient strategy for prey to decrease predation risk than shifting to neighbouring safer habitats but where predators are still present. In known DM systems, however, the DM takes prey not only significantly away from visual predators but also to refuge habitats where these predators are less efficient (the predator evasion hypothesis, Hays 2003), so the effects of DM and habitat shift strategies are confounded by a positive correlation between encounter risk and vulnerability. Therefore, current studies on DM cannot fully shed light on the conditions under which DM may emerge, and cannot distinguish between the relative effects of encounter risk and vulnerability in shaping prey responses.

Here, we tested the hypothesis that predictable encounter risk with a primarily nocturnal predator (Schaller 1972) whose distribution is spatially anchored near prey resource patches, combined with the lack of refuge habitat for the prey, led the prey to develop a DM strategy. We focused on the lion (*Panthera leo*) space use behaviour and on the spatial proactive response of plains zebras (*Equus quagga*) in Hwange National Park (hereafter Hwange NP; Zimbabwe). In this ecosystem, artificial waterholes are associated with large, well grazed, open areas (Chamaillé-Jammes 2009a, Courbin et al. 2016), which are rare (<2% of the study area) in this otherwise wooded savannah. Zebras favour these short-grasslands during daytime (Valeix et al. 2009, Courbin et al. 2016) as they provide profitable forage and high visibility. Lions hunt near waterholes at night (Valeix et al. 2010, Davidson et al. 2013, Courbin et al. 2016), and rest in their vicinity during the day (Valeix et al. 2010, Courbin et al. 2016). We therefore predicted that zebras should display DM, coming close to waterholes during the day to forage and drink when lions are inactive and moving away at night to decrease predation risk when lions become active. Our results supported this prediction, and so we then used lion GPS-tracking data to quantify to what extent these night-time displacements decreased the risk of encountering lions for zebras. Vegetation cover is a significant determinant of zebra vulnerability against lion attacks, as increased cover provides more and better ambush opportunities (Caro 2005, Davidson et al. 2012, Loarie et al. 2013). We therefore also evaluated to what extent day/night changes in the selection for vegetation cover types (e.g. grasslands vs. bushlands) modified the risk of encountering lions. Ultimately, our framework allowed us to compare the relative ability of DM and habitat shift strategies in reducing encounter risk with the predator. In summary, here we (1) quantified the effect of a predator-induced DM in a terrestrial system on the likelihood of encountering a predator, (2) distinguished its effects from those related to day/night changes in selection for vegetation types.

## METHODS

### Study site

The study was conducted in Hwange NP, Zimbabwe. The vegetation is typical of dystrophic semi-arid wooded savannahs (average annual rainfall is c. 600 mm), with woodlands and bushlands interspersed with small grassland patches (Chamaillé-Jammes 2006). We focused on two contrasting seasons: the wet season (November to April) and the late dry season (August to October). During the latter, zebras drink at artificial waterholes (hereafter referred to as ‘waterholes’) that are the only perennial sources of water. All statistical analyses were conducted for both seasons.

In the study area, zebra and lion densities were estimated at c. 100 zebra/100km^2^ (Chamaillé-Jammes 2009b) and c. 3.5 lions/100 km^2^ (Loveridge, unpublished data). Lions hunt for zebras (c. 10% of their diet) but also for other prey (Davidson et al. 2013). The zebra population in Hwange NP seems to be under top-down control by lions, their main predator, and is currently declining due to a high predation pressure (Grange et al. 2015).

### Testing for the existence of zebra DM

We used GPS locations collected hourly from 25 adult female zebras (18 zebras in dry season and 24 zebras in wet season), collared in different herds between August 2009 and November 2013, to assess if zebras systematically moved further away from waterholes at night and if this depended on how close they were to waterholes during the day. For each day and night of each season, we estimated the distance to the closest waterhole (hereafter ‘distance to waterhole’) at which an individual was as the median distance to waterhole over its GPS locations for the given day or night. See Appendix S1: Fig. S1, for the sunrise/sunset-based definition of day/night periods.

We used least-squares spectral fitting to test that distance to waterhole displayed a 24h-periodicity. For each zebra in each season, we visually inspected Lomb-Scargle periodograms (Ruf 1999) for peaks around 24h, and tested the significance of the largest peak within the 20 to 28h window using the randomization procedure implemented in the *lomb* package (Ruf 1999) for the R software (R Development Core Team 2016).

For individuals displaying a significant 24h-periodicity in distance to waterhole (i.e. those performing a diel migration), we investigated if displacement away from waterholes depended upon their proximity to waterholes during the previous day. We did this by modelling the relationship between the night-time distance to waterhole and the distance to waterhole during the previous day using a generalized additive mixed model (GAMM) with thin plate regression splines (Wood 2003). Individuals were included as random factors to account for the unbalanced sampling among individuals (Gillies et al. 2006). The model was fitted using the *gamm4* package (Wood and Scheipl 2014) for the R software (R Development Core Team 2016).

### Characterizing the spatial risk of lion encounter

We first used GPS data from lions, collected over the same period as for zebras, and inhomogeneous point process models (Aarts et al. 2013, Johnson et al. 2013) to build predictive maps of the intensity of lion occurrence within the landscape (see Appendix S2). Separate maps were built for daytime and night-time because lions displayed day/night changes in habitat selection (see Appendix S2: Table S2 and Fig. S1).

We then used these maps to estimate how the risk of encountering lions decreased with the distance to waterhole at zebra locations during daytime (using daytime lion occurrence map and daytime zebra GPS locations; thereafter LionRisk_Day_ZebraUse_Day_ model) and night-time (using night-time lion occurrence map and night-time zebra GPS locations; thereafter LionRisk_Night_ZebraUse_Night_ model). We did so by fitting log-linear mixed-effects models (log-LMM) with the intensity of lion occurrence as the response variable and the log-transformed distance to waterhole as unique predictor, both measured at each zebra GPS location. We allowed for a random intercept for each zebra.

### Quantifying the impact of vegetation types on encounter risk

At any distance to waterhole, the difference between the LionRisk_Day_ZebraUse_Day_ and LionRisk_Night_ZebraUse_Night_ models measured how the combined effects of lion and zebra day/night changes in space use affected their likelihood of encounter. Previous studies conducted in the study area have shown that lions shift to selecting more open vegetation at night (Courbin et al. 2016, see also Appendix S2), and that zebras, although they still select open vegetation types at night, do so less than during the day (Courbin et al. 2016). We disentangled the relative contribution of lion and zebra selection for vegetation types on encounter probability by creating a LionRisk_Night_ZebraUse_Day_ model, fitted on night-time lion occurrence map and daytime zebra GPS locations models. At any distance to waterhole, the difference between this LionRisk_Night_ZebraUse_Day_ model and the LionRisk_Day_ZebraUse_Day_ model measured to what extent lion changes in selection for vegetation types at night would increase zebra risk of encountering lions if zebras behaved as they did during the day. Similarly, the difference between the LionRisk_Night_ZebraUse_Day_ and the LionRisk_Night_ZebraUse_Night_ model measured to what extent zebra changes in selection for vegetation types at night reduced their risk of encountering lions, assuming night-time behaviour for lions. We additionally quantified the overall effect of zebra selection for specific vegetation type on the risk of encountering lions at night by comparing the LionRisk_Night_ZebraUse_Night_ model with one estimated using the night-time lion occurrence map but zebra locations randomly drawn in the landscape (LionRisk_Night_ZebraUse_Random_ model).

### Measuring the effect of DM on encounter risk

Finally, using the above-described models relating the likelihood of encountering lions with the distance to waterhole, we quantified to what extent night-time behavioural adjustments (including both DM and change in selection for vegetation types) allowed zebras to decrease their risk of encountering lions. We first used the GAMM model fitted in section *Testing for the existence of zebra DM* to predict zebra night-time displacement away from waterholes across the range of daytime distances to waterhole. We then used this distance and results from the LionRisk_Night_ZebraUse_Night_ model to predict its night-time risk of encountering lions across a range of distance to waterhole. We then calculated the difference between this risk and the one expected if the zebra did not adjust its behaviour at night (estimating risk at the daytime distance to waterhole from the LionRisk_Night_ZebraUse_Day_ model) across a range of distance to waterhole. To measure the effect of DM only, we compared the decrease in risk brought by all the night-time behavioural adjustments (calculated above from the LionRisk_Night_ZebraUse_Night_ model) with the decrease in risk induced by only the adjustment of the selection for vegetation types. We estimated this latter measure by calculating the difference between the LionRisk_Night_ZebraUse_Night_ and the LionRisk_Night_ZebraUse_Day_ models across a range of distances to waterhole.

## RESULTS

### Zebras undertake DM

During the dry season, zebras were generally within a few km of waterholes, but were closer to waterholes during the day than at night (Figs 1A, 2). Periodogram analyses confirmed that distance to waterhole displayed a well-marked 24h cycle that was significant for 83% of the individuals, while DM frequency varied among individuals (note the variability in normalized power values, Fig. 3A). Zebras moved towards waterholes in the first hours of the morning and moved away at sunset with an average DM of 0.5 +/-0.4 km (mean +/-SD) (see Appendix S1: Fig. S1). However, for zebras with a significant DM pattern, night-time displacement away from waterholes declined as daytime distance from a waterhole increased (Fig. 1B). No night-time displacement away from water occurred beyond a daytime distance of 2.4 km.

**Figure 1.**
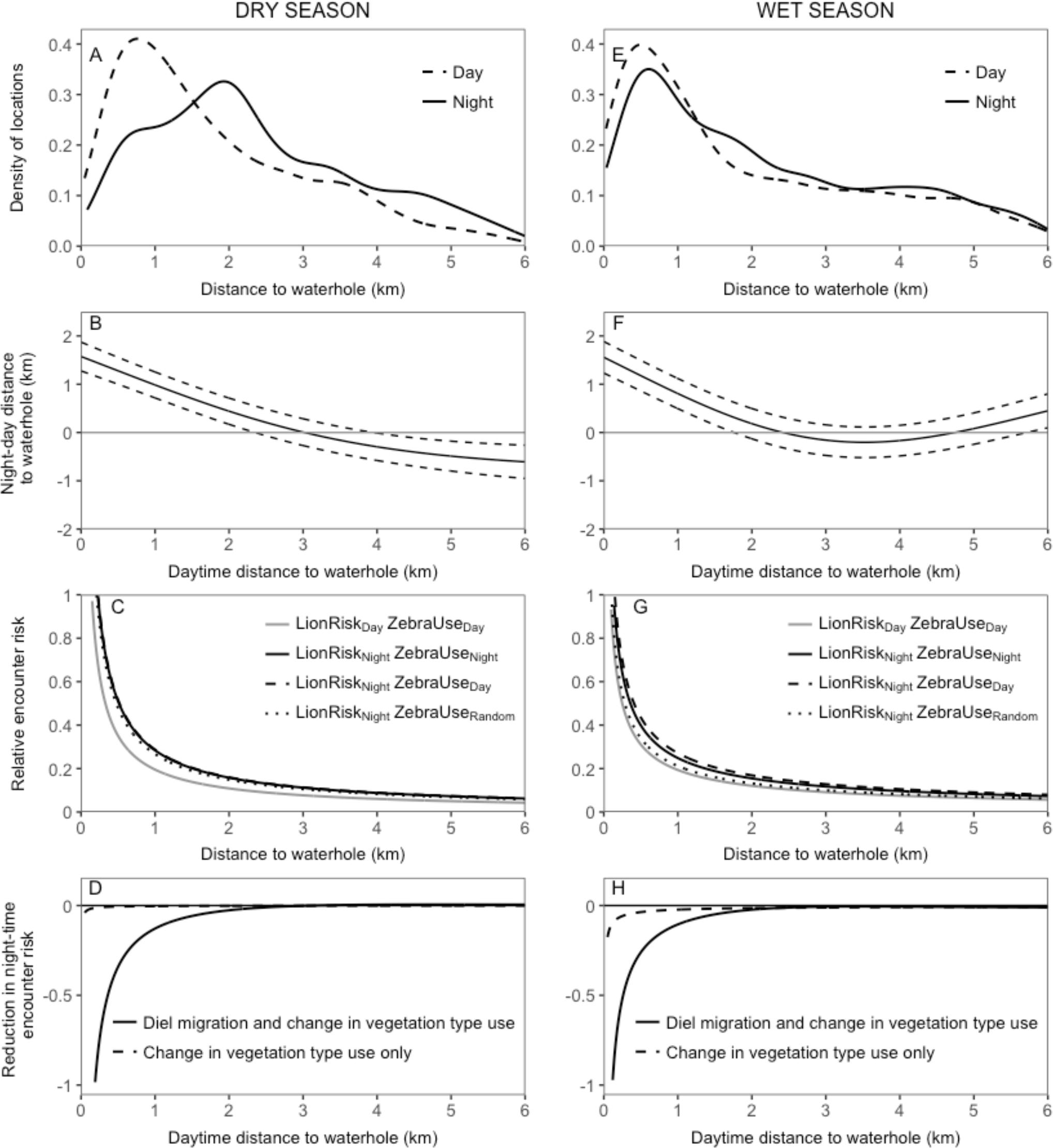
(A,E) Distribution of zebra locations as a function of distance to waterhole during daytime and night-time. The distribution is truncated at 6km (dry season: for both day and night 90% of data are shown and the tail reaches c. 15km; wet season: 76% and 78% of daytime and night-time data are shown, respectively, the tails of the distribution reach 38km [daytime] and 34km [night-time]). (B,F) Difference between zebra distance to waterhole at night and their distance to waterhole during the previous day, as predicted by a generalized additive mixed model (dry season: df=2.827, F=119.8, *P*<0.001; wet season: df=2.986, F=134, *P*<0.001). Positive (negative) values indicate that zebras moved away from (closer to) waterhole at night. Dotted lines represent the 95% confidence interval. (C,G) Expected risk of encountering lions as a function of distance to waterhole and four situations: (1) lion daytime space use and zebra daytime locations (LionRisk_Day_ZebraUse_Day_); (2) lion night-time space use and zebra night-time locations (LionRisk_Night_ZebraUse_Night_), (3) lion night-time space use risk and zebra daytime locations (LionRisk_Night_ZebraUse_Day_), and (4) lion night-time space use risk and random locations (LionRisk_Day_ZebraUse_Random_). (D,H) Differences between the night-time encounter risk expected for zebras displaying full night-time behavioural adjustments (diel migration and change in selection for vegetation types; solid line), or displaying only a change in selection for vegetation types (dotted line), and the night-time risk expected under the assumption of no behavioural adjustment (represented by the horizontal line at 0). More negative values indicated greater reduction in risk.

**Figure 2.**
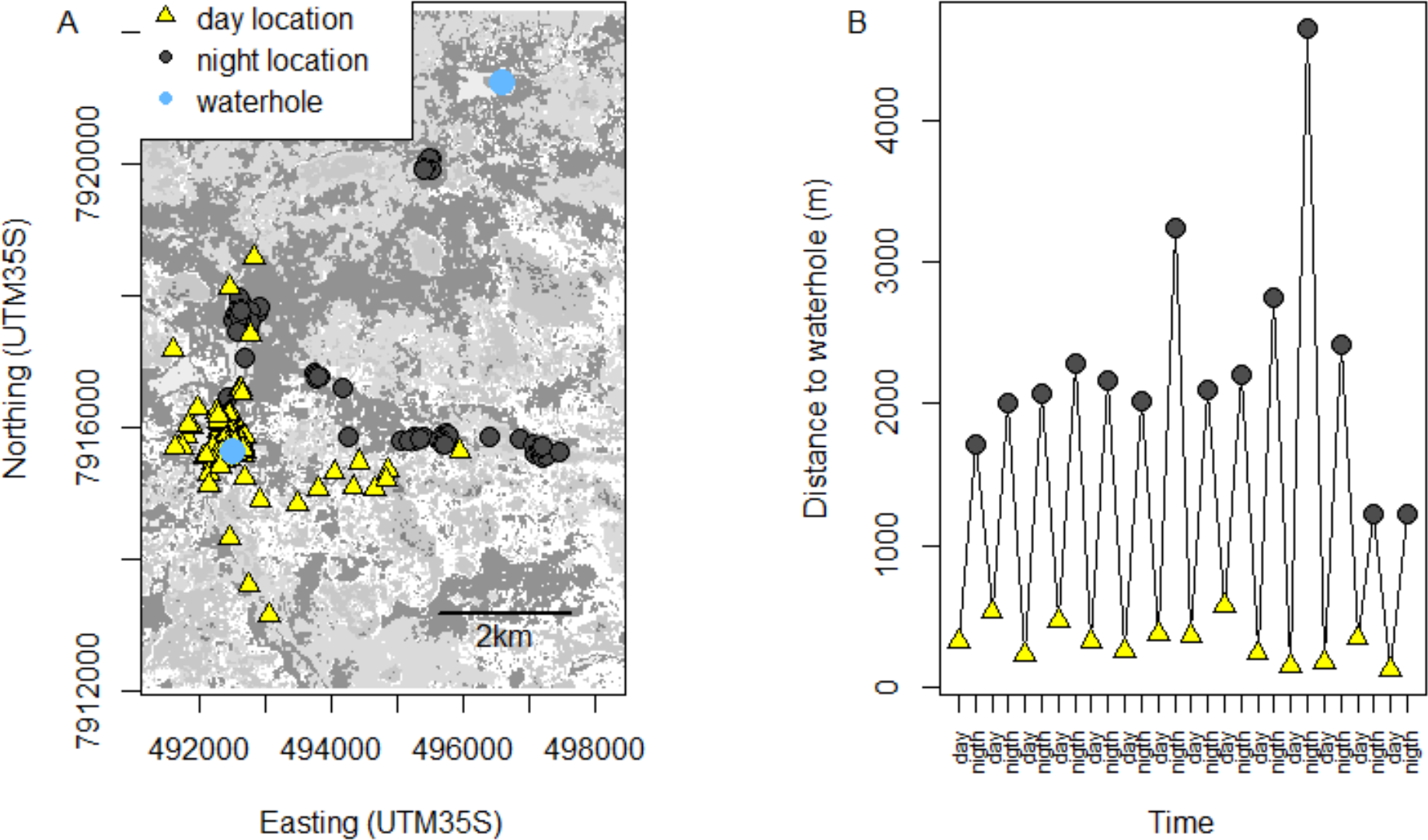
Example of diel migration behaviour. The panels display (A) GPS locations and (B) the median distance to the closest waterhole during day and night, using data obtained from zebra ID AU299 over a 14-day period during the 2009 dry season in the Hwange National Park, Zimbabwe.

**Figure 3.**
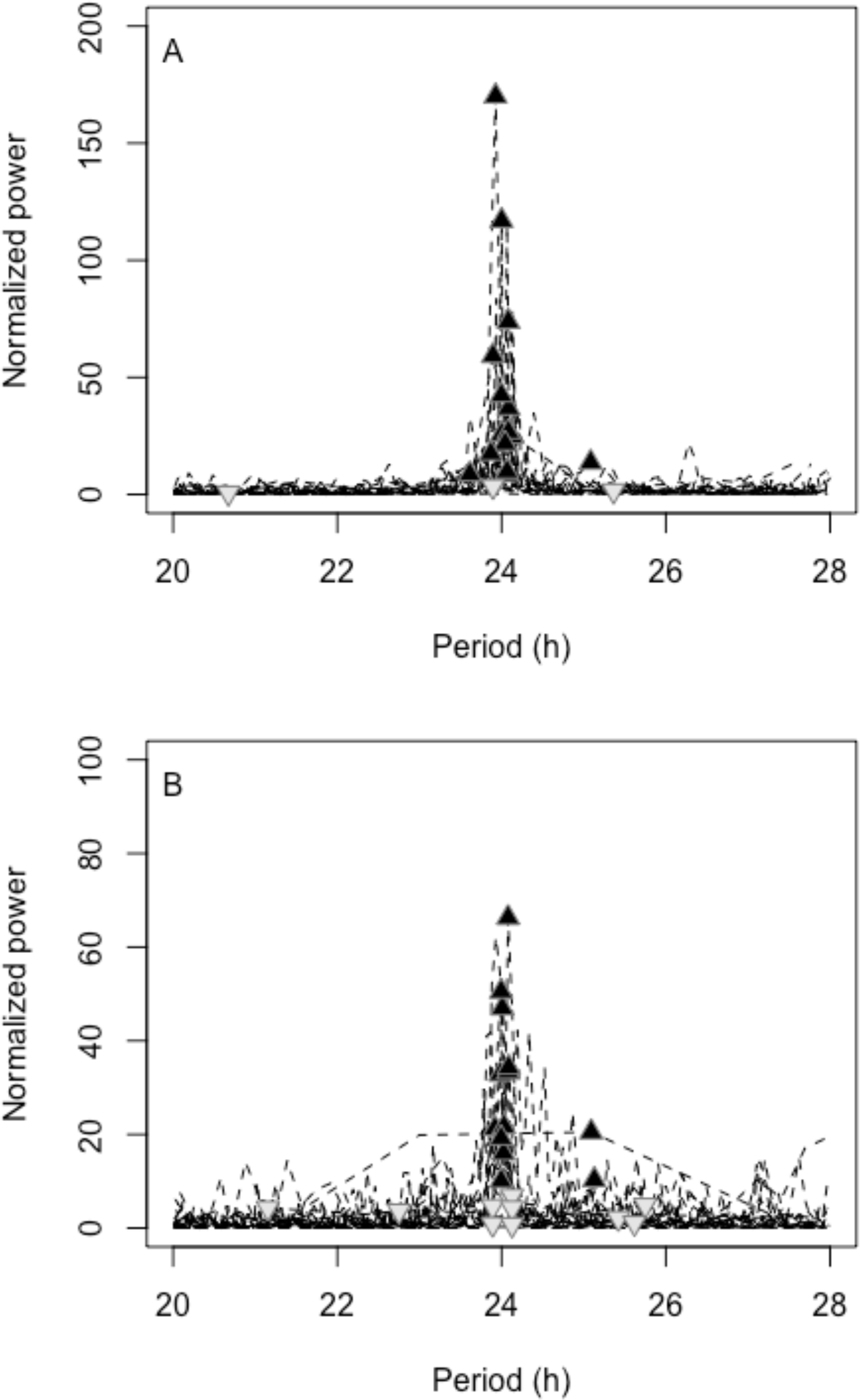
Periodograms of the distance to waterhole time-series for (A) the dry season and (B) the wet season. Each line represents the periodogram for one individual zebra, and the maximum value of each periodogram spectrum within the 20 to 28h-period window is indicated by a triangle. Black triangles pointing up and grey triangles pointing down indicate significant (*P*<0.05) and non-significant (*P*≥0.05) peak values, respectively. Peak values were significant for 83% (15 out of 18) of the individuals in the dry season, and for 54% (13 out of 24) of the individuals in the wet season

During the wet season, zebras remained close to waterholes at night more often than during the dry season (Fig. 1E). Zebras also used DM but, compared to the dry season the 24h-periodicity in back-and-forth movement to waterholes was significant for a lower proportion of zebras (54%) and DMs were less frequent (i.e., lower normalized power values, Fig. 3B). Also, for zebras with a significant DM pattern, the night-time displacements away from waterholes vanished at shorter daytime distance from a waterhole (1.8 km, Fig. 1F).

### Zebra mechanisms of reducing lion encounter risk

During the dry season, zebras’ risk of encountering lions, as indexed by our model, always decreased rapidly with the distance from waterholes (Fig. 1C). At any distance to waterhole, this risk would increase at night if zebras did not adjust their night-time behaviour (Table 1, compare LionRisk_Night_ZebraUse_Day_ and LionRisk_Day_ZebraUse_Day_ models in Fig. 1C). Zebras would be at a higher risk at night because lions always selected for areas close to waterholes, and increased their selection for grasslands and the most open bushlands at night (see Appendix S2: Table S2 and Fig. S1). In response, zebras however only slightly reduced their use of risky vegetation types at night (Table 1, see that LionRisk_Night_ZebraUse_Night_ < LionRisk_Night_ZebraUse_Day_ in Fig. 1C), and these remained highly selected for (Table 1, see that LionRisk_Night_ZebraUse_Night_ > LionRisk_Night_ZebraUse_Random_ in Fig. 1C). Zebras reduced their night-time risk of encountering lions only marginally by decreasing their selection for risky vegetation types (see ‘Night-time risk with changes in vegetation type use only’ in Fig. 1D).

**Table 1.** Log-linear mixed-effect models of the expected risk of encountering lions for zebras as a function of distance to waterhole and four situations: (1) lion daytime space use and zebra daytime locations (LionRisk_Day_ZebraUse_Day_); (2) lion night-time space use and zebra night-time locations (LionRisk_Night_ZebraUse_Night_, serving as baseline), (3) lion night-time space use risk and zebra daytime locations (LionRisk_Night_ZebraUse_Day_), and (4) lion night-time space use risk and random locations (LionRisk_Day_ZebraUse_Random_). Models were fitted for the dry (n=15 individuals) and wet (n=13 individuals) seasons. Coefficients (β) and their 95% confidence intervals (95% CI) are shown. The distance to waterhole was log-transformed.

| Season | Variable | $\beta$ | 95% CI |
| --- | --- | --- | --- |
| Dry | $\text{LionRisk}_{\text{Night}}\text{ZebraUse}_{\text{Night}}$<br>(baseline) | -5.86* | -5.97;-5.76 |
|  | Distance to waterhole | -0.85* | -0.85;-0.84 |
| | $\text{LionRisk}_{\text{Day}}\text{ZebraUse}_{\text{Day}}$ | -0.37* | -0.38;-0.36 |
| | $\text{LionRisk}_{\text{Night}}\text{ZebraUse}_{\text{Day}}$ | 0.01* | 0.002;0.02 |
| | $\text{LionRisk}_{\text{Night}}\text{ZebraUse}_{\text{Random}}$ | -0.06* | -0.07;-0.05 |
| Wet | $\text{LionRisk}_{\text{Night}}\text{ZebraUse}_{\text{Night}}$<br>(baseline) | -6.00* | -6.07;-5.93 |
|  | Distance to waterhole | -0.68* | -0.69;-0.68 |
| | $\text{LionRisk}_{\text{Day}}\text{ZebraUse}_{\text{Day}}$ | -0.25* | -0.26;-0.24 |
| | $\text{LionRisk}_{\text{Night}}\text{ZebraUse}_{\text{Day}}$ | 0.09* | 0.08;0.10 |
| | $\text{LionRisk}_{\text{Night}}\text{ZebraUse}_{\text{Random}}$ | -0.16* | -0.17;-0.15 |

\* 95% confidence intervals exclude zero.

By contrast, the DM that zebras undertook allowed them to dramatically reduce their risk of encountering lions at night (compare curves in Fig. 1D). This reduction was particularly strong when they had spent the daytime near water, where risk would have been high at night (Fig. 1D). Although zebras sometimes stayed at c. 0.6 km away from waterholes at night (Fig. 1A, see Appendix S1: Fig. S2), greater displacements were actually more common (Fig. 1A). Within a few km from waterholes, even moderate displacements had a dramatic influence on the risk of encountering lions: for instance in the dry season, zebras moving from 0.4km to 1.74km away from water between day and night (diel migration = 1.34km, Fig. 1B) decreased by 72% the night-time risk of encountering lions (Fig. 1D). Beyond 2 km, displacement away from water brought little reduction in risk as it was already very low (Fig. 1D).

During the wet season, similar patterns were apparent (Figs 1F-G-H), but because DM where of shorter distances when they occurred (Figs 1E-F), the reduction in encounter risk was lower. Zebras moving from 0.4km to 1.65km away from water between day and night (diel migration = 1.25km, Fig. 1F) decreased by 64% the night-time risk of encountering lions (Fig. 1H).

## DISCUSSION

Our study shows that, in Hwange NP, zebras use areas located near artificial waterholes during daytime, benefiting from the large open grasslands (Chamaillé-Jammes 2009a, Courbin et al. 2016), but routinely moved away from them at risky night period, thus reducing their risk of encountering lions. The diel cycle of predator-avoidance revealed here relies on diel migration and is independent of vegetation cover types, conveying an alternative strategy to the well-known day/night habitat selection shift reported so far in terrestrial systems (Mysterud and Østbye 1999, Kronfeld-Schor and Dayan 2003, Laundré 2010). Indeed, at night zebras still use open grasslands. We did not have data to test whether grass quality is higher closer or further away from waterholes. However, if forage was of better quality away from waterholes, we would expect zebras to forage away from waterholes during the day, as they would find the best resources and be the least likely to be predated upon. This is not what we observed, even less so in the wet season when zebras don’t need to drink and would have foraged away from waterholes during the day if resources were of better quality there. Moreover, the comparison of previous studies suggests that vigilance levels is likely higher when zebras are very near waterholes (Périquet et al. 2010, 2012). Therefore, we assume that predator avoidance, rather than attraction towards better resources, is driving the observed diel migration. Interestingly, the diel migration occurring here or in aquatic systems (Iwasa 1982, Burks et al. 2002, Hays 2003, Benoit-Bird and Au 2006) where no absolute refuge areas occurred differs from the diel response showed by hunted ungulates that take refuge during the day in protected areas completely free of risk (no hunting) (e.g. wild boars [*Sus scrofa* L.] [Tolon et al. 2009] and bison [*Bison bison*] [Fortin et al. 2015]).

### Diel migration is advantageous when space use of the predator is predictable

Diel migration may emerge as an efficient strategy to deal with food-safety trade-offs when prey can reliably identify and travel to places where the absence of a predator is likely (Iwasa 1982, Sainmont et al. 2013). In Hwange NP, lions remain near waterholes most of the time, despite being free to move anywhere (Valeix et al. 2010, Courbin et al. 2016, this study). Areas away from waterholes are therefore predictably safer, and our results show that zebras benefit from this predictability. Zebras have developed a DM strategy allowing them to more than halve their risk of encountering lions during their hunting period, compared to staying to near waterholes. Thus, daily zebra movements to and from waterholes may provide a mechanistic explanation for the low night-time lion-zebra encounter rate observed in Hwange NP (one encounter every 35 days on average, Courbin et al. 2016).

The predictability of the predator distribution however depends on both the landscape configuration and the predators hunting strategies. Ambush sites for sit-and-wait predators are usually predictable (Schmidt and Kuijper 2015), and the actual presence of the predator will be even more predictable if ambush predators focus around prey hotspots. In Hwange NP, the large patches of grasslands located near these waterholes attract grazers and mixed-feeders all year round, and the many water-dependent species naturally use these waterholes during the dry season. Waterholes can thus be considered as prey hotspots in this ecosystem. Lions, being generalist predators (Davidson et al. 2013), may not need to track zebras moving away from waterhole areas if some other prey species do not perform DM. This is yet unknown, but field observations suggest that certain species (e.g. impala, kudu) indeed do not perform DM (unpublished data). This is to be expected as DM should emerge only when predation risk is predictable in space and time, and negatively correlated to resource abundance/quality. This will not be the case for many prey species, especially those that are significantly predated upon by cursorial predators that roam over vast areas and whose distribution is unpredictable (Latombe et al. 2014) Prey of cursorial predators should therefore shift towards safer neighbouring habitats when the predator is detected or is likely to revisit the area rather than moving towards areas where predation risk is uncertain (see examples with wolves, [Creel et al. 2005, Middleton et al. 2013, Latombe et al. 2014, Basille et al. 2015, Schmidt and Kuijper 2015]).

Overall, a better understanding of the landscape and behavioural constraints driving the spatial behaviour of the species making the food web would shed light on the interaction driving the emergence, or not, of diel migration. In this context, it would prove valuable to test the existence of DM in other prey and other ecosystems, contrasting situations with varying levels of prey and predator predictability.

### Does the absence of safe vegetation types facilitate the emergence of DM?

We found that zebras did not alter their selection for vegetation types at night to an extent that would reduce encounter risk with lions significantly. We suspect that this is due to a trade-off between encounter risk and vulnerability across vegetation types. At night, lions strongly select for more open vegetation, possibly to benefit from increased visibility and to maximize encounter rates with prey (Courbin et al. 2016, see Appendix S2: Fig. S1). Zebras could reduce the risk of encountering lions by selecting for more bushy vegetation (see Appendix S2: Fig. S1), but they would then become highly vulnerable in case of an encounter with lions which are primarily ambush/stalking predators (Caro 2005, Davidson et al. 2012, Loarie et al. 2013). Therefore, zebras may decrease encounter risk while maintaining a low vulnerability by conducting DM towards open vegetation types localized in relatively safe areas (i.e. far from waterholes).

### Do DMs have population-level consequences?

Our results could suggest that DM, which strongly decreases zebra likelihood of encountering lions, is a prime determinant of zebra survival rate. However, data from both lion kill surveys (Davidson et al. 2013) and zebra demographic monitoring (Grange et al. 2015) show that adult zebras are less likely to be predated upon by lions during the wet season, when we found that DMs were much less prevalent than in the dry season. It is yet unknown if this seasonal difference in DM patterns is driven by resources or predation sensitivity. They could be linked to the higher cost of leaving the best foraging patches at a time when grass quality is high. Also, it could be that lion favour other prey during the wet season. This itself could be because prey abundance and vulnerability vary seasonally. Adult zebras may be a challenging prey when body condition is good during the wet season (pers. obs.), and this itself could lead them to accept higher chances of meeting lions, especially to forage on good quality grass. All these explanations could explain the lack of relationship between predation rate on adults and prevalence of the diel migration across seasons. Also, this may be because in the wet season lions favour hunting juvenile zebras. Almost half of the juvenile zebras are killed during their first 6 months, mostly by lions (Grange et al. 2015). Therefore, the link between DM and adult predation rate may be distorted by the seasonal presence of juveniles, which may itself constrain the DM as young ones will be less mobile. It remains to be investigated if individual variability in juvenile survival rate could be linked to the ability of some herds to perform longer DM earlier after the birth season. This would allow assessing the population-level consequences of DM, which may occur via consumptive or non-consumptive (e.g. increased energetic expenditure) effects.

## CONCLUSIONS

The study of DM may thus help to clarify the respective roles of encounter risk and vulnerability in driving anti-predation behaviour. Our study emphasizes that DM could possibly be a more general anti-predator strategy than previously thought, and opens new research avenues to better understand the conditions under which it may evolve. In particular, it offers opportunities to study how the behaviour of the predator (i.e. mobility and hunting mode), the constraints for the prey (i.e. resource needs, presence of young) and the spatial context of their interactions (i.e. availability and spatial arrangement of the resource patches) determine the efficiency of DM compared to other anti-predator strategies. Generally, our study answers previous calls to consider the temporal patterns in the predator-prey space race (Hammond et al. 2007). Prey may use high risk, rich food patches during periods of predator inactivity or inefficiency, and move away from these patches when an encounter with the predator becomes more likely or dangerous.

## ACKNOWLEDGMENTS

This research was partly funded by the grants ANR-08-BLAN-0022, ANR-11-CEPS-003, ANR-16-CE02-0001-01, Darwin Initiative for Biodiversity Grants 162-09-015 and EIDPO002. Beyond academic grants, this study was partly funded by Mitsubishi Fund for Europe and Africa, R.G. Frankenberg Foundation, Boesak and Kruger Foundation, Rufford Maurice Laing Foundation, SATIB Trust, Eppley Foundation, Riv and Joan Winant and Recanati-Kaplan Foundation. M. Muzamba, J. Hunt, B. Stapelkamp, N. Elliot and H. Valls Fox contributed to fieldwork. The late N. Ganzin obtained and did some initial work on the Landsat images used for the vegetation map. This research was authorized by the Zimbabwe Parks and Wildlife Management Authority under permits (permit numbers: REF:DM/Gen/(T) 23(1)(c)(ii): 03/2009, 01/2010, 25/2010, 05/2011, 06/2011, 12/2012, 15/2012, 08/2013).

